# Breed strongly predicts hemotropic coinfections in cattle: evidence from a Colombian multispecies molecular survey

**DOI:** 10.1101/2025.06.26.661883

**Authors:** Carolina Ríos Úsuga, Lina María Rendón Ramos, Ingrid Lorena Jaramillo Delgado, Nathalia M Correa-Valencia

**Affiliations:** Research Group of Infectious Diseases, Zoonoses and Environment TestMol Laboratory (GIZMOL), TESTMOL© S.A.S. – Specialized Diagnostic Center, Calle 9C Sur No. 50ff-116, Medellín, Antioquia, Colombia; CENTAURO Research Group, School of Veterinary Medicine, Faculty of Agricultural Sciences, Universidad de Antioquia (UdeA), Calle 70 No. 52-21, Medellín, Antioquia, Colombia

**Author notes:** (NM Correa-Valencia).

## Abstract

Hemotropic pathogen species of *Anaplasma, Babesia, Mycoplasma*, and *Trypanosoma* are endemic to cattle populations and may cause coinfections, complicating disease dynamics and control efforts. However, the host-related factors associated with these infections remain poorly understood under tropical field conditions. This study aimed to investigate the occurrence of hemotropic monoinfections and coinfections in cattle and to identify host-related factors associated with coinfection under field conditions in Colombia. A total of 104 animals were included in the study: 91 cattle, 10 buffaloes, and 3 goats. Among the cattle, 34 (37.4%) exhibited hemotropic monoinfections, 47 (51.6%) were coinfected, and 10 tested negative. Among the buffaloes, 7 (70.0%) presented monoinfections, and 2 (20.0%) were coinfected; among the goats, 2 had monoinfections, and 1 was coinfected— most frequently involving *Mycoplasma* spp. The predominant coinfection patterns were *Anaplasma* + *Mycoplasma* and *Mycoplasma* + *Trypanosoma*, particularly in *Bos indicus* cattle. Bivariate and multivariate analyses identified breed as the strongest predictor of coinfection, with animals of less common breeds showing a 98% lower risk (aOR = 0.02; p = 0.007). Additionally, younger cattle (7–12 months) and *Bos taurus* breed individuals presented reduced odds of coinfection. Our findings reveal a high frequency of hemotropic coinfections in cattle, particularly those involving *Mycoplasma* spp., and demonstrate that host-related factors such as breed and age significantly influence infection risk. *Bos indicus* cattle presented the highest coinfection rates, whereas less common breeds, such as *Bos taurus*, and younger animals presented lower odds. These results highlight the need for surveillance and prevention strategies tailored to the demographic and genetic profiles of livestock populations.

## Introduction

Vector-borne diseases (VBDs) are among the main clinical alterations that generate economic losses due to their morbidity and mortality in animal production. These diseases are distributed throughout the world, from the polar circle to the equator (Demessie et al., 2015). The frequency of VBD is relatively high in tropical and subtropical regions, where its distribution is determined by a complex set of environmental, biological, demographic and social factors (WHO, 2024). Among the countries of Latin America and the Caribbean, Colombia is considered one of the ten countries with the highest risk of VBD, since approximately 85% of the territory is located below 1,600 m.a.s.l., where climatic, geographic and epidemiological conditions favor the transmission of these pathogens (Carrasquilla et al., 2023).

In Colombia, stable to unstable enzootic conditions with different prevalence rates may predispose cattle to cycles of transmission or continuous outbreaks when infected individuals are present (Ferreira et al., 2022). In extensive and semiextensive livestock production, where strategies to improve management practices under changing climatic conditions are limited, the proliferation of hematophagous arthropods and the persistence of pathogens such as *Babesia bovis, B. bigemina, Anaplasma marginale, Trypanosoma vivax* and *T. evansi* in single or mixed infections are favored (Murcia-Mono et al., 2025). In South American countries, such as Colombia, the variation in prevalence levels, including stable or unstable enzootic endemism, predisposes individuals to enzootic cycles or epizootic outbreaks in the presence of infectious cattle (Ferreira et al., 2022; Mor et al., 2024), which makes the diagnosis, management and control of these diseases difficult.

According to Resolution 003714 of the Colombian Agricultural Institute (ICA), diseases of national interest in cattle include babesiosis, anaplasmosis and trypanosomiasis because of their significant impact on health and productivity (ICA, 2015). Effective therapeutic and prophylactic monitoring of these hemotropic agents requires rapid, reliable and highly sensitive diagnostic methods. Common conventional methods, such as blood smears and serology, cover the basic needs for diagnosis but have disadvantages in terms of sensitivity and specificity. Serodiagnosis does not differ between current and past infections (Salih et al., 2015). On the other hand, in endemic regions, due to high exposure, molecular amplification tests, such as PCR, allow the identification of infectious agents at levels well below the detection limit of commonly used techniques (Oliveira-Sequeira et al., 2005).

To date, there are few epidemiological studies of vector-borne diseases in cattle and buffaloes, and studies in small ruminants are lacking. In addition, the intrinsic and extrinsic factors associated with infection are not known. In addition, the frequency of Hemoplasmas in ruminants in Colombia is unknown. Therefore, this study aimed to investigate the occurrence of hemotropic monoinfections and coinfections in cattle, buffaloes, and goats and to identify host-related factors associated with coinfection under field conditions in Colombia.

## Materials and methods

### Ethical considerations

No animals were manipulated during the study, as data from routine molecular diagnoses of infectious diseases in ruminant samples sent to the TestMol© S.A.S laboratory were analyzed. The use of each animal’s data for research was authorized through a sample submission form signed by those in charge of requesting the tests. Therefore, ethics committee approval for animal experimentation was not needed.

### Study design and study population

A cross-sectional study was carried out, considering records of bovines, buffaloes, and goats that tested positive by real-time PCR for hemotropic agents (*Anaplasma* spp., *Babesia* spp., *Mycoplasma* spp., and *Trypanosoma* spp.) from samples (i.e., blood, fecal matter, tissues, and nasal secretions) submitted by veterinarians from herds, farms or organizations for livestock production in different regions of Colombia (consisting of eight provinces) to TestMol© S.A.S., a molecular reference laboratory, throughout the year 2024 (January 1^st^ to December 31^st^). All the samples from the farm animals were sent to the laboratory for molecular diagnosis to detect hemotropic agents. Based on the obtained molecular results (hemotropic profiles), every animal was evaluated for monoinfection and coinfection with the other agents considered in the study. Demographic variables such as species, breed, sex, and age group were collected from the sample submission form used to request sample shipping and processing at TestMol© S.A.S. This manuscript was prepared in accordance with the Strengthening the Reporting of Observational Studies in Epidemiology (STROBE) guidelines for cross-sectional studies.

### Genomic DNA isolation from blood samples

The DNA from bacteria and parasites was extracted from a 200 μL blood sample. Samples from 104 individuals were extracted via an automatic method with Kingfisher™ Duo open extraction equipment (Thermo Fisher Scientific Inc., MA, USA) and a MagMAX™ CORE M Express-96 nucleic acid purification kit (Thermo Fisher Scientific, Waltham, MA, USA) according to the established conditions of the manufacturer. The final elution volume of the samples at the end of the extraction process was 100 μL. The DNA was stored at -80°C until processing. The purity and concentration of the DNA were analyzed with a Nanodrop spectrophotometer (NANO-400, A&E Lab., GuanDong, China). Samples with a DNA concentration greater than 10 ng/µl and a purity ratio of 260/280 between 1.8 and 2.0 were considered suitable for processing via qPCR. The samples were re-extracted if they did not meet the criteria until they passed the required quality parameters.

### qPCR method for identification of *T. gondii* and hemotropic agents

The DNA samples were processed with specific primers (Macrogen, Korea) targeting the 16S rRNA gene for bacterial pathogens (*Anaplasma* spp. and *Mycoplasma* spp.) and the 18S rRNA gene for protozoan parasites (*Babesia* spp. and *Trypanosoma* spp.). For the amplification process, reactions of 20 μl were prepared containing 10 μl of BlasTaq™ Green (2X) qPCR MasterMix enzyme (Applied Biological Materials Inc. (abm), BC, Canada), 1 μl of each primer (100 mM), 3 μl of DNA from each sample (20–25 ng/μl) and 5 μl of ultrapure water. The thermal profile was run with an initial denaturation of 3 minutes at 94°C, then 35 cycles of 10 seconds at 95°C, 15 seconds at 57°C, and 10 seconds at 72°C, followed by a final extension for 1 minute at 72°C. The simple qPCRs for each infectious agent were performed in a Mic qPCR Cycler with 4 channels (BioMolecular Systems, Australia) according to the laboratory’s protocols.

This study used both positive and negative controls to ensure the accuracy and validity of the results. The positive controls were provided by TestMol© S.A.S., which consists of a qPCR mixture with the corresponding primers plus the plasmid for each evaluated infectious agent in the same volume as the sample DNA used in a reaction. As a negative control, DNase-RNase-free sterile water (Cat No. 129114; Qiagen, Germany) was used. For the internal controls used for DNA extraction and qPCR, specific primers for the cytochrome B gene in mammals were used (Pfeiffer et al., 2004).

The results were analyzed in terms of the reached threshold cycle (Ct) value reached and the dissociation curves (temperature range Tm(°C) established for each infectious agent). Samples with a Ct value <35 were considered positive, and samples with values ≥35 were considered negative. The bacterial and parasitic loads in copies/μl were calculated based on the laboratory’s internal test validation process, where the theoretical copy number for each plasmid was established.

### Data analysis

All the data collected during the study were compiled in Microsoft Excel spreadsheets (Microsoft Corp., Redmond, WA, USA) and exported to Stata v.18 (StataCorp, 2023, College Station, TX, USA) for statistical analysis and data visualization. Demographic and contextual variables were categorized for analytical purposes. Age was grouped into four biologically relevant categories: Group 1 (calves) from 0 to 6 months, Group 2 (juveniles) from 7 to 12 months, Group 3 (heifers) from 13 to 24 months, and Group 4 (adults) ≥25 months of age. Breeds were classified as either purebred or mixed breed, while sex was recorded as male or female. Geographic origin was determined based on the municipality and province of sampling. Coinfection status, the primary outcome, was defined as the simultaneous molecular detection of two or more hemotropic pathogens (*Anaplasma* spp., *Babesia* spp., *Mycoplasma* spp., and *Trypanosoma* spp.) and dichotomized as present or absent. Descriptive statistics were used to summarize the distributions of the demographic and diagnostic variables. To assess associations between coinfection status and each independent variable, bivariate analyses were performed via Fisher’s exact test or the chiquare (χ^2^) test, depending on expected cell frequencies. Fisher’s exact test was used when any expected cell count was less than five, particularly in 2×2 tables, whereas the χ^2^ test was applied when all expected counts met or exceeded this threshold. A significance level of p<0.05 was established for all inferential analyses. To further evaluate the independent contribution of each variable to coinfection risk and to control for potential confounding factors, a multivariate logistic regression model was developed. Variables yielding p<0.20 in the bivariate analysis were considered for inclusion. Stepwise backward elimination was used to refine the model, and adjusted odds ratios (aORs) with corresponding 95% confidence intervals (CIs) were reported. Multicollinearity was assessed via variance inflation factors (VIFs), and model fit was evaluated via the Hosmer–Lemeshow goodness-of-fit test and the Bayesian information criterion (BIC). Residual diagnostics and influence statistics were also examined to ensure model robustness. A heatmap was generated to illustrate the distribution of coinfections, facilitating the identification of potential epidemiological patterns within the bovine study population.

## Results

A total of 104 animals were included in the study: 91 cattle, 10 buffaloes, and 3 goats. A geographic distribution map was generated for all animal species of interest according to hemotropic pathogen-positive cases and detected agents (Fig 1). Please note that if the province or the categorized breed data were missing or if a mixed breed record was found, it was not considered in the map.

**Fig 1.**
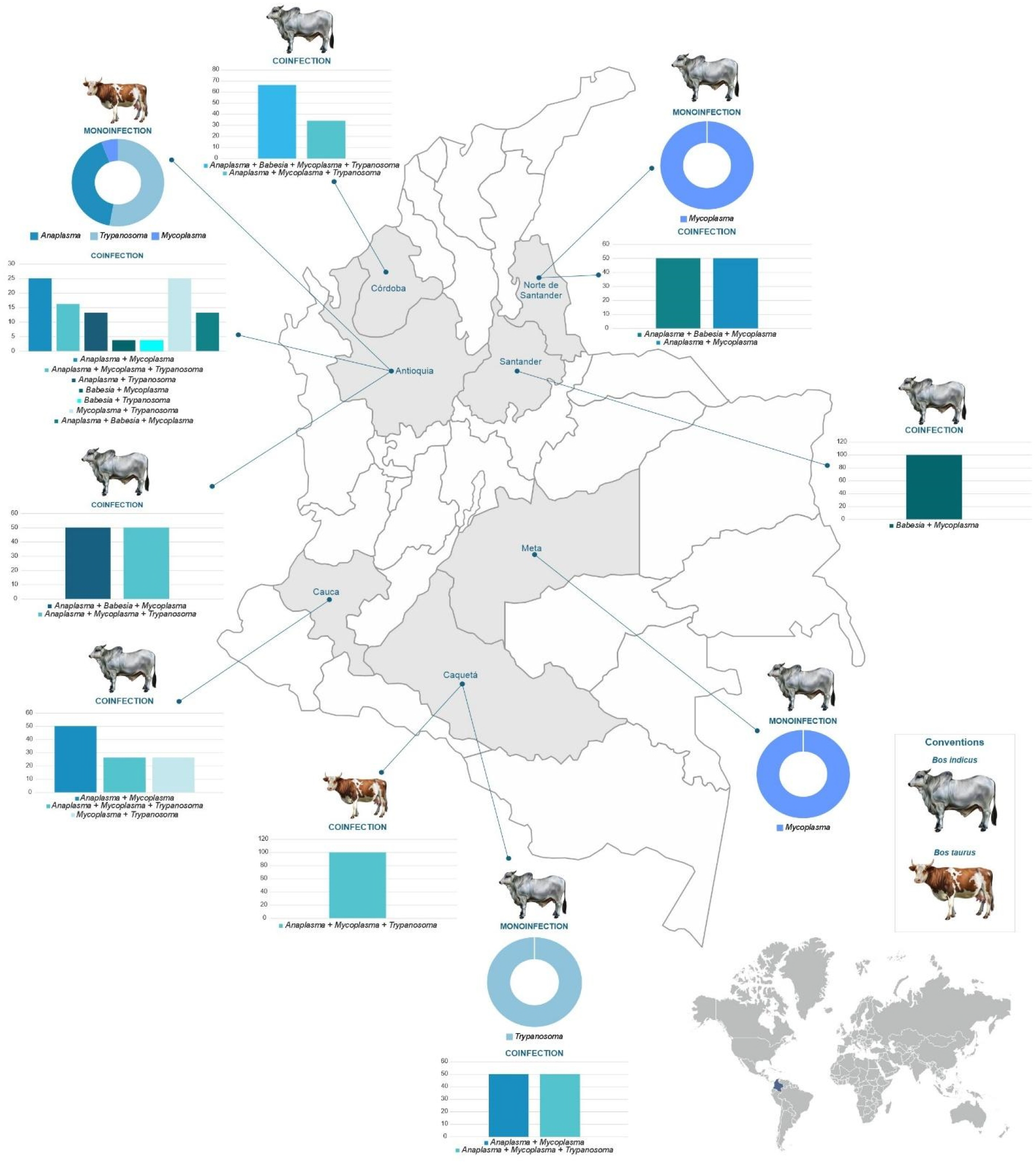
Geographic distribution of hemotropic pathogen-positive cases by animal species and detected agents in the bovine population (mono- and coinfections), 2024, Colombia.

Among the cattle, 34 (37.4%) had hemotropic monoinfections, 47 (51.6%) were coinfected, and 10 tested negative. Table 1 presents a descriptive summary of the bovine population evaluated in this study. The variables included demographic and management-related characteristics such as age group, sex, breed, geographic origin and infection status.

**Table 1.** Descriptive characteristics and hemotropic infection status of the bovine population included in the study (n = 91), 2024, Colombia.

| Variable | Category | n |
| --- | --- | --- |
| Province | Antioquia | 48 |
|  | Caquetá | 26 |
|  | Cauca | 4 |
|  | Chocó | 0 |
|  | Córdoba | 4 |
|  | Meta | 3 |
|  | Norte de Santander | 3 |
|  | Not reported | 2 |
|  | Santander | 1 |
| Breed | Brahman | 12 |
|  | Zebu | 3 |
|  | Mixed breed | 7 |
|  | Gyr | 7 |
|  | Hereford | 1 |
|  | Holstein | 29 |
|  | Jersey × Holstein | 11 |
|  | Jersey | 3 |
|  | Not reported | 16 |
|  | Swiss brown | 1 |
|  | Red sindhi | 1 |
| Categorized breed* | *Bos indicus* | 22 |
|  | *Bos taurus* | 46 |
|  | Mixed breed | 7 |
|  | Not reported | 16 |
| Sex* | Female | 57 |
|  | Male | 29 |
|  | Not reported | 5 |
| Age group* | 1 (calves) | 15 |
|  | 2 (juvenile) | 6 |
|  | 3 (heifers) | 28 |
|  | 4 (adults) | 35 |
|  | Not reported | 7 |
| *Anaplasma* spp. | Negative | 47 |
|  | Positive | 44 |
| *Babesia* spp. | Negative | 76 |
|  | Positive | 15 |
| *Mycoplasma* spp. | Negative | 30 |
|  | Positive | 61 |
| *Trypanosoma* spp. | Negative | 63 |
|  | Positive | 28 |
| Monoinfection | No | 57 |
|  | Yes | 34 |
| Coinfection | No | 44 |
|  | Yes | 47 |

Table 2 displays the results of the bivariate analysis assessing associations between selected independent variables, such as breed, categorized breed, sex, and age group, and the detection of each hemotropic pathogen (*Anaplasma, Babesia, Mycoplasma*, and *Trypanosoma*) independently. Only animals with monoinfection were considered for this analysis to avoid potential confounding by coinfection status. Bivariate analyses revealed significant associations between several host-related variables and the molecular detection of specific hemotropic pathogens among monoinfected cattle. Breed was significantly associated with the detection of *Anaplasma* (p = 0.024) and *Trypanosoma* (p = 0.036). When breeds were categorized, significant associations were observed with *Anaplasma* (p = 0.012), *Mycoplasma* (p = 0.035), and *Trypanosoma* (p = 0.046). Additionally, sex was significantly associated with *Anaplasma* detection (p = 0.025).

**Table 2.**
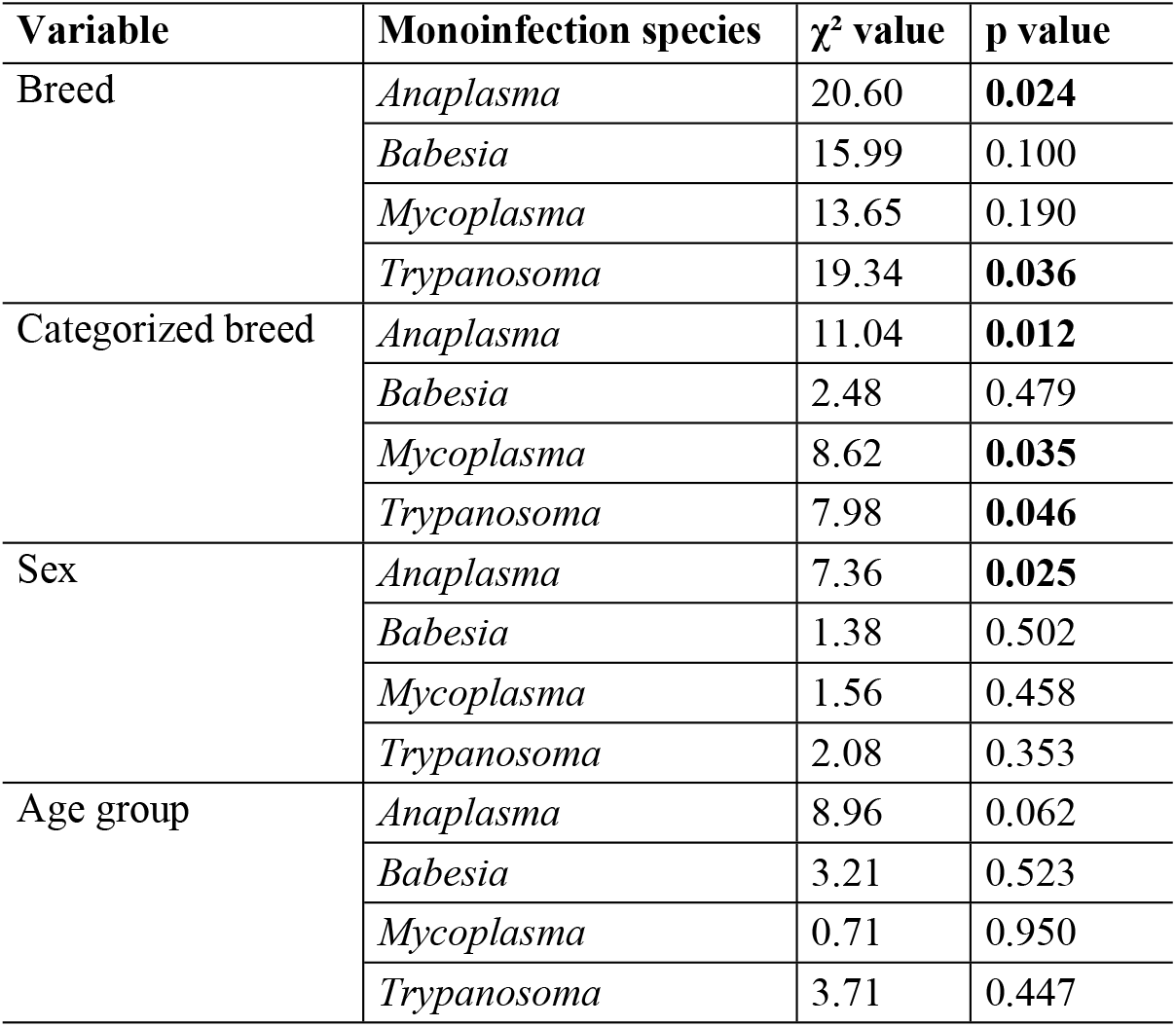
Bivariate associations between host-related variables and hemotropic pathogen detection in the bovine population included in the study (n = 91), 2024, Colombia.

Among the 91 cattle included in the study, several host-related variables were significantly associated with coinfection status. Significant associations were found for breed (p = 0.007), categorized breed (p = 0.002), sex (p = 0.026), and age group (p = 0.041). Following the general analysis of coinfection status, each coinfection type was analyzed separately in relation to host-related variables, including breed, categorized breed, sex, and age group. This disaggregated analysis aims to identify differential associations that may be masked when considering coinfection as a single aggregated outcome. Notably, significant associations were observed between host-related factors and specific hemotropic coinfection patterns in cattle. For example, *Anaplasma* + *Mycoplasma* and *Mycoplasma* + *Trypanosoma* coinfections were strongly associated with breed and categorized breed, suggesting potential genetic or physiological predispositions. The combination of *Anaplasma* + *Mycoplasma* was also significantly associated with sex, whereas *Anaplasma* + *Trypanosoma* was strongly related to breed. Moreover, the triple coinfection *Anaplasma* + *Mycoplasma* + *Trypanosoma* was significantly associated with both breed (p = 0.001) and categorized breed (p = 0.016), reinforcing the influence of genetic background on susceptibility. Although no significant associations were found with sex or age group, a marginal trend was observed for age, suggesting that immunological maturity or early-life exposure may also play a role. Table 3 presents a detailed overview of the bivariate associations for each coinfection profile.

**Table 3.** Bivariate associations between host-related variables and specific hemotropic coinfection types by pathogen species in the bovine population included in the study (n = 91), 2024, Colombia.

| Variable | Coinfection species | $\chi^2$ value | p value |
| --- | --- | --- | --- |
| Breed | *Anaplasma* + *Babesia* | 13.378 | 0.342 |
| Categorized breed | *Anaplasma* + *Babesia* | 3.519 | 0.318 |
| Sex | *Anaplasma* + *Babesia* | 0.509 | 0.775 |
| Age group | *Anaplasma* + *Babesia* | 5.316 | 0.256 |
| Breed | *Anaplasma* + *Mycoplasma* | 33.096 | **0.001** |
| Categorized breed | *Anaplasma* + *Mycoplasma* | 15.771 | **0.001** |
| Sex | *Anaplasma* + *Mycoplasma* | 6.368 | **0.041** |
| Age group | *Anaplasma* + *Mycoplasma* | 11.57 | **0.021** |
| Breed | *Anaplasma* + *Trypanosoma* | 27.136 | **0.007** |
| Categorized breed | *Anaplasma* + *Trypanosoma* | 8.325 | **0.04** |
| Sex | *Anaplasma* + *Trypanosoma* | 5.084 | 0.079 |
| Age group | *Anaplasma* + *Trypanosoma* | 10.501 | 0.033 |
| Breed | *Babesia + Mycoplasma* | 24.318 | **0.018** |
| Categorized breed | *Babesia + Mycoplasma* | 3.807 | 0.283 |
| Sex | *Babesia + Mycoplasma* | 0.685 | 0.71 |
| Age group | *Babesia + Mycoplasma* | 2.919 | 0.572 |
| Breed | *Babesia + Trypanosoma* | 7.376 | 0.832 |
| Categorized breed | *Babesia + Trypanosoma* | 2.147 | 0.542 |
| Sex | *Babesia + Trypanosoma* | 0.347 | 0.841 |
| Age group | *Babesia + Trypanosoma* | 3.566 | 0.468 |
| Breed | *Mycoplasma + Trypanosoma* | 28.192 | **0.005** |
| Categorized breed | *Mycoplasma + Trypanosoma* | 8.611 | **0.035** |
| Sex | *Mycoplasma + Trypanosoma* | 3.056 | 0.217 |
| Age group | *Mycoplasma + Trypanosoma* | 3.132 | 0.536 |
| Breed | *Anaplasma + Babesia + Mycoplasma* | 16.492 | 0.17 |
| Categorized breed | *Anaplasma + Babesia + Mycoplasma* | 5.415 | 0.144 |
| Sex | *Anaplasma + Babesia + Mycoplasma* | 0.459 | 0.795 |
| Age group | *Anaplasma + Babesia + Mycoplasma* | 6.604 | 0.158 |
| Breed | *Anaplasma + Babesia + Trypanosoma* | 15.634 | 0.209 |
| Categorized breed | *Anaplasma + Babesia + Trypanosoma* | 6.414 | 0.093 |
| Sex | *Anaplasma + Babesia + Trypanosoma* | 1.174 | 0.556 |
| Age group | *Anaplasma + Babesia + Trypanosoma* | 3.852 | 0.426 |
| Breed | *Anaplasma + Mycoplasma + Trypanosoma* | 31.937 | **0.001** |
| Categorized breed | *Anaplasma + Mycoplasma + Trypanosoma* | 10.38 | **0.016** |
| Sex | *Anaplasma + Mycoplasma + Trypanosoma* | 3.535 | 0.171 |
| Age group | *Anaplasma + Mycoplasma + Trypanosoma* | 7.864 | 0.097 |
| Breed | *Babesia + Mycoplasma + Trypanosoma* | 10.909 | 0.537 |
| Categorized breed | *Babesia + Mycoplasma + Trypanosoma* | 4.561 | 0.207 |
| Sex | *Babesia + Mycoplasma + Trypanosoma* | 0.156 | 0.925 |
| Age group | *Babesia + Mycoplasma + Trypanosoma* | 2.152 | 0.708 |
| Breed | *Anaplasma + Babesia + Mycoplasma + Trypanosoma* | 15.634 | 0.209 |
| Categorized breed | *Anaplasma + Babesia + Mycoplasma + Trypanosoma* | 6.414 | 0.093 |
| Sex | *Anaplasma + Babesia + Mycoplasma + Trypanosoma* | 1.174 | 0.556 |
| Age group | *Anaplasma + Babesia + Mycoplasma + Trypanosoma* | 3.852 | 0.426 |

A multivariate logistic regression model was developed to evaluate the independent contributions of the selected variables to the risk of coinfection in bovines (Table 4). The variables included in the model were selected based on biological relevance and bivariate associations. The aORs, 95% CIs, and corresponding p values are reported. The model included sex, age group, simplified categorized breed, and simplified province of origin, based on bivariate results with p < 0.20. Categories with low frequency were grouped to ensure model stability. The final model converged successfully and revealed aORs with 95% CIs. All the predictors had a VIF below 5, indicating no significant multicollinearity. The highest VIF was observed for the intercept, which is not of concern. The BIC indicated a good model fit, and the direction and magnitude of effects were consistent with prior descriptive trends. The variable with the strongest independent association with coinfection was the breed classification. Animals categorized as belonging to less common breeds (“Other”) were significantly less likely to present coinfections, with an aOR of 0.02 (p = 0.007), indicating a 98% reduction in odds compared with the reference group. The age group also had a notable effect: cattle aged 7 to 12 months had an aOR of 0.08 (p = 0.061), suggesting a 92% lower likelihood of coinfection relative to adult animals, although this result was near the threshold of statistical significance. A similar trend was observed for animals identified as *Bos taurus*, with an aOR of 0.05 (p = 0.08), indicating a lower probability of coinfection than the reference breed category. Animals for which age data were not recorded also presented a decreased risk (aOR = 0.15; p = 0.08). The intercept of the model reflects the baseline log odds of coinfection when all the predictors are at their reference levels.

**Table 4.** Multivariate logistic regression model for factors associated with hemotropic coinfection in the bovine population included in the study (n = 91), 2024, Colombia.

| Model term (predictor) | Coefficient | p value | Adjusted Odds Ratio (aOR) |
| --- | --- | --- | --- |
| Breed category: Other* | -3.69 | 0.00 | 0.02 |
| Intercept | 4.25 | 0.02 | 70.35 |
| Age group: 7–12 months** | -2.56 | 0.06 | 0.08 |
| Breed category: *Bos taurus* ** | -2.97 | 0.08 | 0.05 |
| Age group: Not reported** | -1.91 | 0.08 | 0.15 |
| Age group: >25 months | -1.14 | 0.13 | 0.32 |
| Department: Other | -2.49 | 0.17 | 0.08 |
| Sex: Male | -0.75 | 0.36 | 0.47 |
| Sex: Not reported | 1.18 | 0.48 | 3.27 |
| Province: Caquetá | -0.9 | 0.53 | 0.4 |
| Age group: 13–24 months | -0.2 | 0.85 | 0.82 |

\*Statistically significant ( $p < 0.05$ ), \*\*Marginally significant ( $0.05 \leq p < 0.10$ )

Fig 2 displays the proportion of individuals within each demographic subgroup —defined by sex, age group, and categorized breed, that were simultaneously positive for pairs of pathogens.

**Fig 2.**
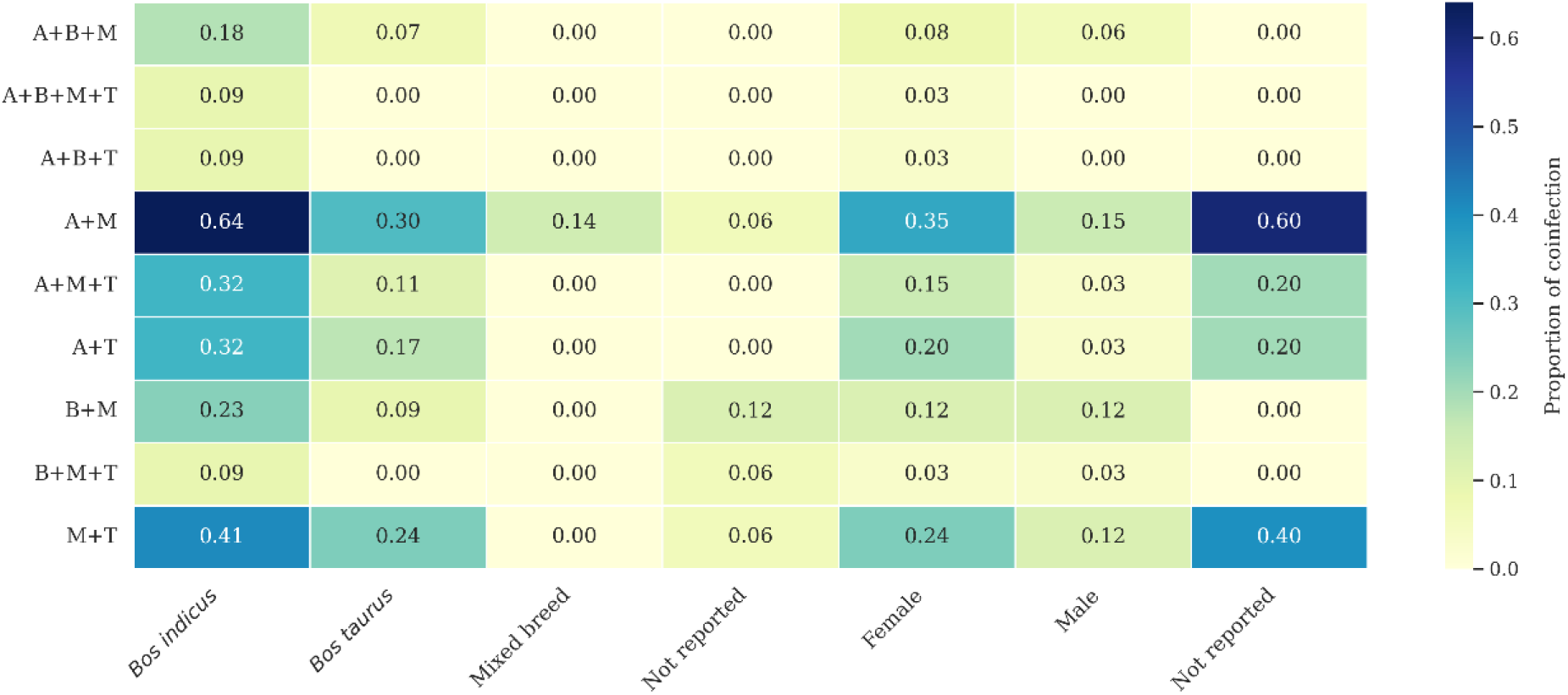
Heatmap of coinfection proportions by demographic subgroup (categorized breed and sex) in the bovine population included in the study (n = 91), 2024, Colombia. A=*Anaplasma*; B=*Babesia*; M=*Mycoplasma*; T=*Trypanosoma*. Darker shades indicate a greater proportion of coinfection within that specific subgroup.

The province of Caquetá was the most common sampling location, accounting for half of the records of the 11 buffalo specimens included in the study. The most frequently detected pathogen was *Mycoplasma* spp., which was found in all infected cases. Seven buffaloes were monoinfected with *Mycoplasma* spp., while two individuals presented coinfections: one with *Mycoplasma* + *Trypanosoma* and another with *Mycoplasma* + *Babesia*. Only one buffalo tested negative for all pathogens.

All three goats originated from the municipality of Medellín (province of Antioquia). Among the three goat samples analyzed, hemotropic infection was confirmed in two individuals. One animal was monoinfected with *Mycoplasma* spp., while the other was coinfected with *Anaplasma* + *Mycoplasma*. None of the goats were positive for *Babesia* spp. or *Trypanosoma* spp.

## Discussion

In Colombia, vector-borne diseases are among the most important problems in animal production. In the Andean region of southwestern Colombia, there is evidence of a prevalence of *Anaplasma* spp. (40.3%), *Babesia* spp. (41.9%) and *Trypanosoma* spp. (45%) in cattle (Murcia-Mono., 2025). Another molecular study in northeastern Colombia reported 83.7%, 59.9% and 40.9% positivity for *B. bigemina, A. marginale* and *B. bovis*, respectively (Jaimes-Dueñez., 2024). In our study, we found that, in cattle, the frequencies of infection with *Anaplasma* spp., *Babesia* spp. and *Trypanosoma* spp. were 48.3%, 16.5% and 30.8%, respectively.

In buffaloes, a molecular prevalence of 6.5% for *B. bigemina* and 17.7% for *B. bovis* has been reported in two regions of Colombia (Jaimes-Dueñez, 2018). With respect to small ruminants (sheep and goats), only one study is known in Valledupar, Colombia, where DNA of *Coxiella burneti* was found in 6% of sheep and 0.6% of goats (Contreras et al., 2018). In the present study, monoinfection and coinfection were observed in buffaloes and goats, with *Mycoplasma* spp. being the main infectious agent identified.

This is the first report of the molecular detection of *Mycoplasma* spp. in cattle, buffaloes and goats. Therefore, these results provide new insights into the epidemiology of hemotropic pathogens in cattle, buffalo and goats in Colombia. By combining molecular detection methods with detailed host-related data, we were able to identify key demographic and genetic factors associated with the occurrence of both monoinfections and coinfections. These findings have important implications for understanding pathogen ecology and for specific surveillance and control strategies in livestock populations.

Among the 104 animals examined, 45.2% were coinfected with two or more hemotropic agents. In cattle, coinfections were present at 51.6%, and the main coinfection detected was *Anaplasma* spp. + *Mycoplasma* spp., followed by *Anaplasma* spp. + *Trypanosoma* spp. and *Mycoplasma* spp. + *Babesia* spp. Coinfections of *B. divergens* + *A. Phagocytophilum* in *Bos taurus* cattle have been reported in the literature previously (Andersson et al., 2017). In addition, coinfections with different species of *Anaplasma* spp. and *Erhlichia* spp. have also been reported (André et al., 2020; Miranda et al., 2021).

In buffaloes, we detected coinfection of *Mycoplasma* spp. + *Trypanosoma* spp. and *Mycoplasma* spp. + *Babesia* spp. in two individuals. There are other reports of coinfection with vector-borne diseases, such as concomitant infection with *A. marginale* + *Theileria annulata* (Fatima et al., 2024) and, likewise, coinfection with two species of *Babesia* spp. (*B. bovis* and *B. bigemina*) and A. *marginale* in water buffaloes (Obregón et al., 2019). Although coinfections are more common in cattle than in buffaloes and goats, these findings highlight the importance of evaluating the interactions of multiple pathogens in production animals and their economic impact.

When performing bivariate analysis, we observed that certain host-related variables were significantly associated with the detection of specific hemotropic pathogens in the study cattle. Breed was significantly associated with *Anaplasma* spp. and *Trypanosoma* spp. positivity. Similarly, the categorized breed also showed a significant association with *Anaplasma* spp., *Mycoplasma* spp., and *Trypanosoma* spp. infection. Another study also revealed that the breed variable had a significant effect on the presence of *Theileria mutans* in cattle (Moumouni et al., 2018). However, an in vivo study revealed that *B. indicus* and *B. taurus* are equally susceptible to *T. parva* infection (Ndungu et al., 2005). Nevertheless, more studies with other hemotropic agents in endemic areas such as Colombia are lacking.

Additionally, sex was associated with *Anaplasma* detection, as has been reported in other studies, where sex was a significant determinant of *Anaplasma* spp. infection (Kispotta et al., 2016; Noaman., 2020). No significant associations were found between age groups and any of the pathogens analyzed independently, suggesting that age may not be a major determinant of infection status in this bovine population. Similarly, *Babesia* infection was not significantly associated with any of the host-related variables evaluated. These findings indicate that, among cattle, factors such as breed, categorized breed, and sex appear to play a more prominent role in susceptibility to *Anaplasma, Trypanosoma*, and *Mycoplasma* infections.

When coinfection was considered as a single outcome, statistically significant associations were observed for breed, categorized breed, sex, and age group in cattle. No other studies in the literature have shown that breed and sex are variables associated with infection with various hemotropic agents. However, one study reported that dual infection (*T. evansi* + *A. marginale*) occurred 2.5 times in calves (Sharma et al., 2015). These results suggest that both genetic background and demographic factors may influence susceptibility to coinfection.

The observed associations between specific coinfection patterns and host-related variables underscore the potential role of genetic and developmental factors in modulating susceptibility to hemotropic pathogens. The consistent links between coinfections, such as *Anaplasma* spp. *+ Mycoplasma* spp., *Mycoplasma* spp. *+ Trypanosoma* spp., and *Anaplasma* spp. *+ Trypanosoma* spp., and both breed and categorized breeds suggest that breed-specific immunogenetic traits may influence infection dynamics. A situation that has been previously reported, where purebred *Bos indicus* cattle are relatively more resistant to *Babesia. bovis* (Bock et al., 1997). *Bos Taurus* cattle appear to be more likely to develop severe, acute anaplasmic disease than *Bos indicus* cattle are (Abdisa, 2019).

The disproportionately high coinfection rates among *Bos indicus* animals further supports this hypothesis. Moreover, the increased occurrence of certain coinfections in younger animals suggests the possible contribution of early-life exposure or an immature immune response. The transplacental transmission of these hemotropic agents should also be considered, since this type of transmission has been previously reported (Da Silva et al., 2014) and should be studied to establish control programs in the country. Notably, the strong associations of triple coinfection with *Anaplasma + Mycoplasma + Trypanosoma* with breed-related variables reinforce the importance of considering genetic background in future risk assessments. Although age and sex did not show statistically significant associations in this case, the nearly significant trend observed for age suggests that temporal immune factors may still warrant further investigation.

In the multivariate model, categorized breed emerged as the strongest independent predictor of coinfection. Compared with those in the reference group, the animals classified under less common breeds (“Other”) were 98% less likely to present coinfection. Age also appeared to play a role: cattle aged 7–12 months had 92% lower odds of coinfection — although this result was just slightly above the typical threshold for statistical significance, and *Bos taurus* animals showed a similar trend with a 95% reduction in odds compared with the reference group. These findings reinforce the relevance of host genetic background and age in shaping coinfection dynamics in the bovine study population.

This study presents limitations that should be considered when interpreting the results. In cattle, while the sample size allowed for robust statistical analyses, some variables included categories with low representation, which may have influenced the precision of certain estimates. In buffaloes, despite a slightly broader geographic representation than in goats, the overall number of individuals sampled remained small. This limited the ability to perform subgroup comparisons or draw strong inferential conclusions for this species. In goats, all samples originated from a single location, resulting in complete homogeneity for the location variable and minimal variation in other demographic characteristics. Additionally, the very limited sample size and low variability in infection status precluded the use of statistical tests such as Fisher’s exact test or chi-square analysis. As a result, no meaningful statistical associations could be assessed. These limitations emphasize the need for future studies involving larger and more demographically diverse populations, particularly goats and buffaloes, to better characterize the epidemiology and risk factors associated with hemotropic infections in these species.

## Conclusion

The results of this study highlight the complexity of hemotropic pathogen dynamics in cattle and the significant influence of host-related factors, particularly breed, age, and sex, on infection and coinfection patterns. A notably high frequency of coinfections was observed, especially involving *Mycoplasma* spp., with *Bos indicus* cattle showing the highest rates, particularly for the *Anaplasma* + *Mycoplasma* and *Mycoplasma* + *Trypanosoma* combinations. In the multivariate model, breed classification emerged as the strongest predictor of coinfection: animals classified as fewer common breeds presented a 98% lower risk. Reduced odds of coinfection were also observed among younger age groups and *Bos taurus* cattle. These findings underscore the importance of accounting for host-specific and local epidemiological factors when designing surveillance systems and preventive strategies. The identification of such associations supports the need for targeted, context-specific approaches to disease monitoring and control. Tailoring interventions to the demographic and genetic profiles of livestock populations may improve the detection, management, and ultimately prevention of vector-borne and blood-borne infections in cattle.

## Author contributions

**Conceptualization:** Carolina Ríos Úsuga, Lina María Rendón Ramos.

**Data curation:** Nathalia M Correa Valencia.

**Formal analysis:** Nathalia M Correa Valencia.

**Methodology:** Carolina Ríos Úsuga, Lina María Rendón Ramos, Ingrid Lorena Jaramillo Delgado, Nathalia M Correa Valencia.

**Writing – original draft:** Carolina Ríos Úsuga, Lina María Rendón Ramos, Nathalia M Correa Valencia.

**Writing – review & editing:** Carolina Ríos Úsuga, Lina María Rendón Ramos, Ingrid Lorena Jaramillo Delgado, Nathalia M Correa Valencia.

## Data availability

The test conditions of the molecular processes are part of the intellectual property of the company TestMol© S.A.S. However, if there is a need for additional data or details, such information can be obtained through a request process that maintains complete confidentiality. Other data will be made available on request.

## Declaration of generative AI and AI-assisted technologies in the writing process

During the preparation of this work the authors used OpenAI ChatGPT (May 2025 version, https://chat.openai.com/) to improve the readability and language of the manuscript. After using this tool, the authors reviewed and edited the content as needed and took full responsibility for the content of the published article.

